# βVPE is involved in tapetal degradation and pollen development by activating proprotease maturation in *Arabidopsis thaliana*

**DOI:** 10.1101/503508

**Authors:** Ziyi Cheng, Bin Yin, Jiaxue Zhang, Yadi Liu, Bing Wang, Hui Li, Hai Lu

## Abstract

Vacuolar processing enzyme (VPE) is responsible for the maturation and activation of vacuolar proteins in plants. We found that *βVPE* was involved in tapetal degradation and pollen development by transforming proproteases into mature protease in *Arabidopsis thaliana. βVPE* was expressed specifically in the tapetum from stages 5-8 of anther development. The *βVPE* protein first appeared as a proenzyme and transformed into the mature enzyme before stages 7-8. The recombinant *βVPE* protein self-cleaved and transformed to a 27-kD mature protein at pH 5.2. The mature *βVPE* protein could induce the maturation of CEP1 *in vitro. βvpe* mutants exhibited delayed vacuolar degradation and decreased pollen fertility. The maturation of CEP1, RD19A, and RD19C were seriously inhibited in *βvpe* mutants. Our results indicate that βVPE is a crucial processing enzyme that directly participates in the maturation of cysteine proteases before vacuolar degradation, and is indirectly involved in pollen development and tapetal cell degradation.

**Highlight:** βVPE is a crucial processing enzyme that directly participates in the maturation of cysteine proteases before vacuolar degradation, and is indirectly involved in pollen development and tapetal cell degradation.

## 1. Introduction

Tapetal cells are degraded through programmed cell death (PCD) to provide various nutrients for pollen development, particularly the formation of pollen exine. Premature or abrogated PCD of tapetal cells can disrupt the supply of these nutrients to microspores, resulting in sterile pollen (Ko *et al.*, 2003; Varnier *et al.*, 2005). Many cysteine protease enzymes are ubiquitously involved in tapetal cell degeneration and pollen development (Song *et al.*, 2016). For example, OsCP1, a rice cysteine protease gene, plays an important role in pollen development and is regulated by PTC1 and TDR (Lee *et al.*, 2004; Lin *et al.*, 2017; Fu *et al.*, 2014). BnMs3 participates in tapetum development, microspore release, and pollen-wall formation in *Brassica napus* (Zhou *et al.*, 2012). Previous studies in our lab revealed that CEP1 plays an irreplaceable executor role during tapetal PCD and affects pollen development (Zhang *et al.*, 2014). A vacuolar system mediated by vacuolar processing enzyme (VPE) is considered a cellular suicide strategy in plant development and cell death programs (Tang *et al.*, 2016; Teper-Bamnolker *et al.*, 2017). VPE mediates the initial activation of some other vacuolar enzymes, which then degrade the vacuolar membrane and initiate the proteolytic cascade leading to PCD (Gong *et al.*, 2018; Hatsugai *et al.*, 2015; Lam, 2005; Yamada *et al.*, 1999). VPE is also capable of processing several seed proteins, including the 2S albumins and 11S globulins (Shimada *et al.*, 2003).

VPEs, which are asparagine-specific cysteine proteinases exclusively located in the vacuoles of plants, are synthesized as larger, inactive proprotein precursors, from which the C-terminal and N-terminal propeptides are sequentially removed in acidic conditions (pH 5.5) via self-catalysis to produce the active mature forms (Kuroyanagi *et al.*, 2002; Misas-Villamil *et al.*, 2013). The Arabidopsis genome contains four VPE genes (αVPE, βVPE, γVPE, and δVPE), which can be separated into two subfamilies: the vegetative-type αVPE and γVPE, and the seed-type βVPE and δVPE (Greenberg and Yao, 2004; Kinoshita *et al.*, 1999; Gruis *et al.*, 2002). Previous research into plant VPEs has mostly focused on plant senescence, terminal differentiation, and pathogen-induced hypersensitive cell death. αVPE and γVPE, which are upregulated during wounding, senescence, and pathogen infection, may play vital roles in plant cell death (Kinoshita *et al.*, 1999; Yamada *et al.*, 2004). Promoter-GUS analyses revealed the up-regulation of αVPE in dying cortex cells located next to the emerging lateral root (Kinoshita *et al.*, 1999) and of γVPE in dying circular-cell clusters of anthers during the later stages of pollen development (Hatsugai, 2015). In contrast, βVPE is essential for storage protein processing (Shimada *et al.*, 2003), and δVPE, which is specifically expressed in the seed coat, is associated with cell death (Nakaune *et al.*, 2005). OsVPE1, which is a homolog of the Arabidopsis βVPE gene, is a cysteine protease that plays a crucial role in the maturation of rice glutelins (Wang *et al.*, 2009). Additionally, abnormal accumulation of the precursors of 12S globulins has been reported in Arabidopsis mutants lacking VPE (Gruis *et al.*, 2002; Shimada *et al.*, 2003).

Previous studies have found that *βVPE* expression is significantly downregulated in *spl/nzz* and *ems1/exs* Arabidopsis mutants, which display dramatically altered anther cell differentiation patterns (Wijeratne *et al.*, 2007). In particular, expression of βVPE is detected in the stamen, flower pedicel, pollen, petal, carpel, and sepal during flower development, as well as in seeds and root tips (Kinoshita *et al.*, 1999). βVPE is expressed in roots, flowers, buds, and ovules, and is specifically expressed during ovule development in *Vitis vinifera* (Tang *et al.*, 2016). For these reasons, βVPE is speculated to play an essential role in flower development, but its exact function and corresponding mechanism of action remain uncertain.

To investigate the role of βVPE in anther development, we characterized the expression of *βVPE* in the anther and the phenotype of the *βvpe* mutant. We also detected the maturation of cysteine proteinases CEP1, RD19A, and RD19C by βVPE *in vitro* and *in vivo*. Our results indicate that βVPE acts as a trigger during anther development by activating cysteine proteinases in acidic vacuolar environments. Here, we present the first evidence of the activation of papain-like cysteine proteases by VPE during anther development in *Arabidopsis thaliana*.

## 2. Methods

### 2.1 Plant materials and growth conditions

*Arabidopsis thaliana* accession Columbia was used as the wild-type control. Plants were grown in a soil mixture (3:1:1 mixture of peat moss-enriched soil:vermiculite:perlite) with a 14-h light/10-h dark photoperiod at 23°C. Homozygous T-DNA insertion mutants were identified by polymerase chain reaction (PCR) using βVPE-F/R/B primers (F: 5’-TGACCAATTCCACAAACTTCC-3’; R: 5’-TGTCGGCATAAGAATCTTTGG-3’; B: 5’-T CAAACAG GATTTTCGCCT GCT-3’).

### 2.2 Characterization of the mutant phenotype

Arabidopsis plants were photographed using a digital camera (Coolpix 9100; Nikon, Tokyo, Japan). Arabidopsis pod and pollen germination images were acquired using an M165 C microscope (Leica, Wetzlar, Germany). The germination rates of mutant and wild-type seeds were measured using 1/2 Murashige and Skoog (MS) culture medium. To evaluate the viability of mature pollen grains, germination was assessed by culturing fresh pollen grains in germination medium (pH 5.8) containing 3 mM CaCl_2_, 1 mM H_3_BO_3_, 56 mM inositol, 1% (w/v) agar, and 15% (w/v) sucrose at 25°C for 3 h. For each group, 200 pollen grains were counted. Each experiment was repeated three times with both mutants and wild-type plants.

### 2.3 Paraffin sections

Freshly dehisced anthers were collected at stages 8-13 from both wild-type and mutant plants and fixed in formaldehyde-acetic acid-ethanol (FAA) for four hours (Zhang et al., 2014). The samples were then prestained with 1% safranin overnight at room temperature before being dehydrated in an alcohol gradient series (1 h each at 70, 85, 90, and 100% alcohol) and cleared in a xylene/alcohol gradient series (1 h each at 70, 85, 90, and 100% xylene). The samples were incubated in xylene/paraffin (1:1) overnight at 38°C and dipped in 58°C paraffin three times (1 h per incubation). Paraffin-embedded samples were sectioned according to the manual of molecular experiments(Han *et al.*, 2018) and observed under an M165 C microscope.

### 2.4 Scanning electron microscopy (SEM)

Pollen grains collected from freshly dehisced anthers of both wild-type and mutant plants were mounted on SEM stubs. The mounted samples were coated with palladium-gold in a sputter coater (E-1010; Hitachi, Tokyo, Japan) and examined by SEM (S-3400N; Hitachi) at an acceleration voltage of 10 kV.

### 2.5 Transmission electron microscopy (TEM)

Both wild-type and mutant anthers at stages 9-12 were prefixed and embedded (Han et al., 2018). Ultrathin sections (70 nm) were obtained with a UC6 ultramicrotome (Leica) and double-stained with 2% (w/v) uranyl acetate and 2.6% (w/v) lead citrate aqueous solution. Observations and image capture were performed with an H-7650 TEM (Hitachi) at 80 kV and an 832 charge-coupled device camera (Gatan, Inc., Pleasanton, CA, USA).

### 2.6 Molecular Cloning and Plasmid Construction

An 1847-bp promoter of *βVPE* (Pro*βVPE*) was amplified with Pro-βVPE-F/R primers (F: 5’-ATAAGTAGTAATATCAAGTTC-3’; R: 5’-C AATTGTCTAATTATATTTAAT-3’). The promoter was cloned into the pCAMBIA 1300 vector for *Proβvpe:GUS* fusion construct and transformed into Arabidopsis. The open reading frame (ORF) minus the first 63 bp of βVPE cDNA was amplified by PCR with the two βVPE-CM-F/R primers (F: 5’-GAGTCACGCGGTCGGTTCGAG-3’; R: 5’TCAGGCGCTATAGCCTAAG-3’) and inserted downstream of the pET30a plasmid T7 promoter (Novagen). The expression, extraction, purification, and renaturation of the CEP1 and βVPE proteins were performed according to the procedure described by Zhang et al. (2014).

### 2.7 qRT-PCR Analyses

*βVPE* expression in different Arabidopsis tissues and buds of different stages was assessed by qRT-PCR using SYBR Green qPCR mix (LabAid; Thermo Scientific) on an iQ5 Multicolor Real-Time PCR detection system (Bio-Rad) using the *qRT-βvpe-F/R* primers (F: 5’-GATTCTTATGCCGACAGAGG-3’; R: 5’-CCTGGTGTCTGTAGTTTCCA-3’). The PCR conditions were as follows: 94°C for 3 min; 40 cycles at 94°C for 10 s, 55°C for 20 s, 72°C for 20 s, and 60°C for 30 s; and 72°C for 1 min. To normalize expression data, qRT-actin-F/R was used as an internal control. Data were analyzed using the iQ5 (Bio-Rad) software, and differences in gene expression were calculated using the 2-DDCt analysis method.

### 2.8 Immunoblotting

Antimature-βVPE antibody at a 1:200 dilution followed by affinity-purified goat anti-rabbit IgG horseradish peroxidase (HRP)-conjugated antibody (CW Bio) at a 1:4000 dilution were used for immunoblotting. ECL Plus protein gel blotting detection reagents (CW Bio) were used as the HRP substrate and were exposed in the Fusion X7 chemiluminescence imaging system.

### 2.9 GUS staining assay

Flower buds at different stages from heterozygous transgenic lines were treated with 90% (v/v) acetone for 0.5 h on ice. The tissues were subsequently stained with X-Gluc solution and incubated at 37°C for 12 h to visualize GUS activity. The samples were cleared with 75% (v/v) ethanol prior to visualization. The treated samples were subsequently embedded in Technovit 7100 resin. Images were captured under an M165 C microscope.

### Accession Numbers

Sequence data from this study can be found in the Arabidopsis Genome Initiative database under accession numbers AT1G62710(βVPE), AT5G50260 (CEP1), AT4G39090 (RD19A), and AT4G16190 (RD19C).

## 3. Result

### 3.1 Characterization of *βVPE* Expression in Arabidopsis

To identify the function of *βVPE* (AT1G62710) during Arabidopsis anther development, we investigated *βVPE* expression characteristics. We performed qRT-PCR analysis with total RNA extracted from various organs, including roots, stems, leaves, and buds. *βVPE* was highly expressed in flower buds but almost undetectable in roots, stems, and leaves (Figure 1A). The development of Arabidopsis anthers is divided into 14 stages based on morphological landmarks that correspond to cellular events visible under a microscope (Sanders *et al.*, 1999). The expression of *βVPE* appeared in stages 5-6, reached its maximum level in stages 7-8, and then declined sharply to a barely detectable level in stages 9-12 (Figure 1B). An 1847-bp promoter of *βVPE* (Pro*βVPE*) was cloned into the pCAMBIA 1300 vector for the *Proβvpe:GUS* fusion construct and transformed into Arabidopsis. GUS activity was detected in the bud, including the stamen, pistils, and sepals (Figure 1C-F). GUS activity was detected in the anther during stages 5-8, declined sharply in stages 9-10, and was almost undetectable in stages 11-13 (Figure 1G-I).

**Figure 1.**
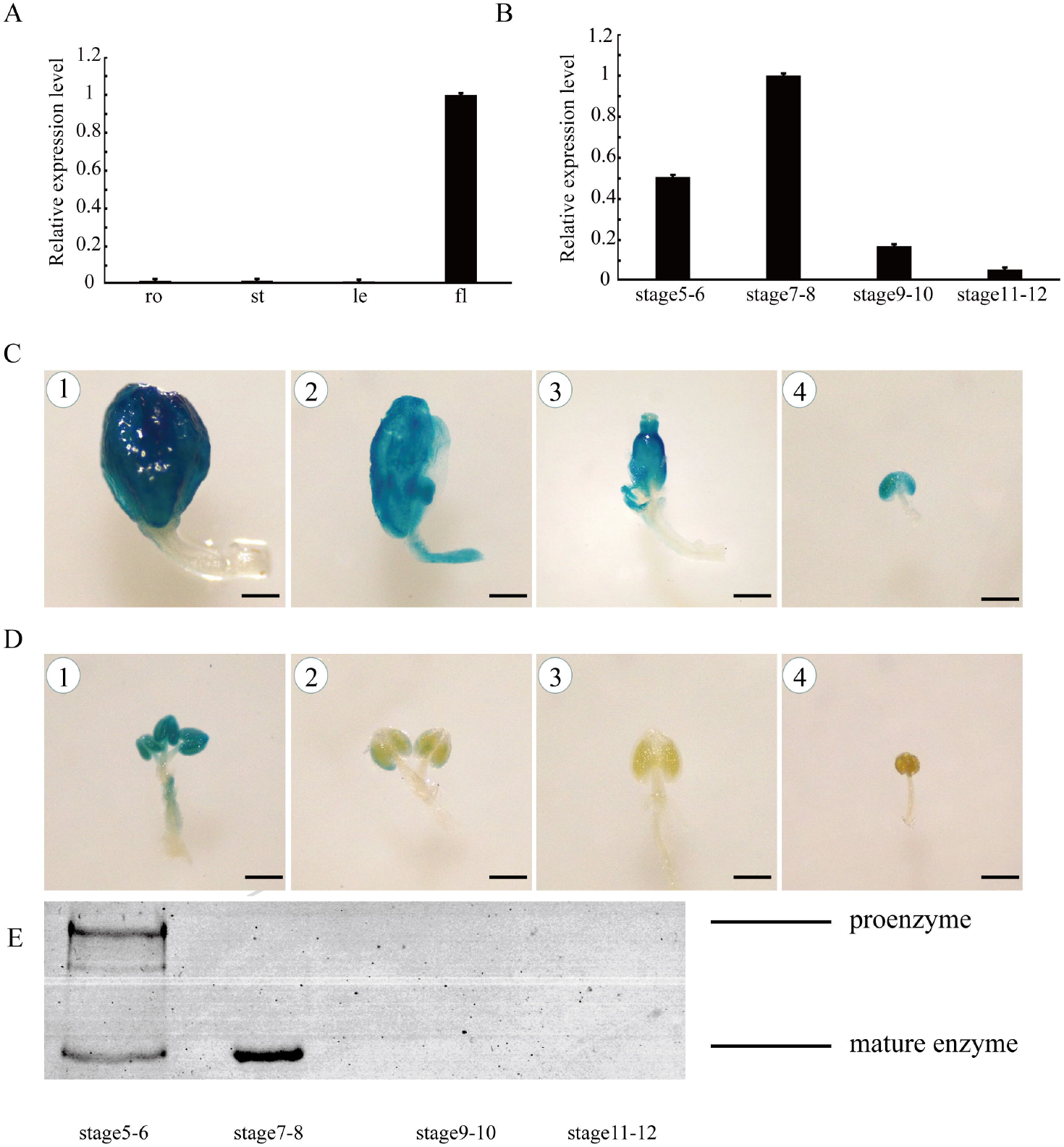
*βVPE* expression pattern (A) *βVPE* spatial and temporal expression analyses performed by qRT-PCR. Fl, flower; le, leaf; ro, root; st, stem. (B) qRT-PCR of *βVPE* expression in wild-type bud tissues at different developmental stages. Bars represent standard deviations. The expression of *βVPE* in stages 7-8 was selected as 1. (C) Histochemical assay for GUS activity harboring the *βVPE* promoter-GUS fusion in bud: (C1) buds; (C2) petal; (C3) pistil; (C4) stamen; (D) GUS activity during anther development: (D1) anther at stage 7-8; (D2) anther at stage 9-10; (D3) anther at stage 11-12; (D4) anther at stage 13-14. (M) Immunoblot analysis of total anther protein extracts from stages 5-12 with antimature-*βVPE* antibody.

We performed immunoblotting using an antimature-βVPE antibody in anthers from stages 5-12 to evaluate βVPE maturation time. The results revealed that only the 51-kD proenzyme was detected in stages 5-6, while the mature 27-kD mature enzyme appeared during stages 5-8. However, during stages 9-12, the quantity of the 51-kD proenzyme and the mature 27-kD enzyme greatly decreased, becoming barely detectable (Figure 6M).

Taken together, these results indicate that the *βVPE* gene is expressed abundantly in Arabidopsis anther development from stages 5-8. First, the premature protease is expressed at stages 5-6, and then the proenzyme is transformed to the mature form with acidification of the vacuole during stages 5-8. After stage 8, *βVPE* is barely expressed and the mature βVPE protein rapidly degrades.

### 3.2 Morphology of βvpe mutants

To identify the function of *βVPE* during *Arabidopsis* anther development, we obtained two T-DNA insertion mutants (CS_1007412 and SAIL_50_F12) from the Arabidopsis Biological Resource Center (ABRC). The T-DNA was inserted into the promoter of *βVPE* in the two mutants (Figure 2A and B). The *βVPE* expression levels in mutant buds were analyzed by qRT-PCR (Figure 2C). The Arabidopsis *ACTIN1* gene (AT2G37620) was used as the reference for normalization. The results revealed that the expression of *βVPE* was completely suppressed in the mutants. In both lines, plants displayed reduced male fertility, and normal female development was confirmed by reciprocal cross analysis. Given that the two mutants both displayed reduced male fertility, CS_1007412 was used for further analysis and is referred to as *βvpe-1* hereafter. *βvpe-1* mutant plants displayed a normal (wild-type) phenotype during vegetative and early generative development stages. The germination rate of pollen grains *in vitro* was significantly lower in *βvpe-1* (45.13%, 116 of 257; P<0.05) than in the wild type (85.54%, 219 of 256; P<0.05; Figure 2D and E). A scanning electron microscopy examination revealed that mature pollen grains in wild-type plants were uniformly spheroid and had finely reticulate ornamentation on their surfaces (Figure 2F and H), while the *βvpe-1* mutant produced some mature pollen grains that were similar to wild-type and some abnormal pollen grains (50.00%, 56 of 112, P<0.05) exhibiting shrunken, collapsed, and gemmate-baculate sculpture without regularly reticulate ornamentation (Figure 2G and I). These results indicate that the absence of *βVPE* markedly impaired pollen development and resulted in sterile pollen grains with abnormal pollen morphology.

**Figure 2.**
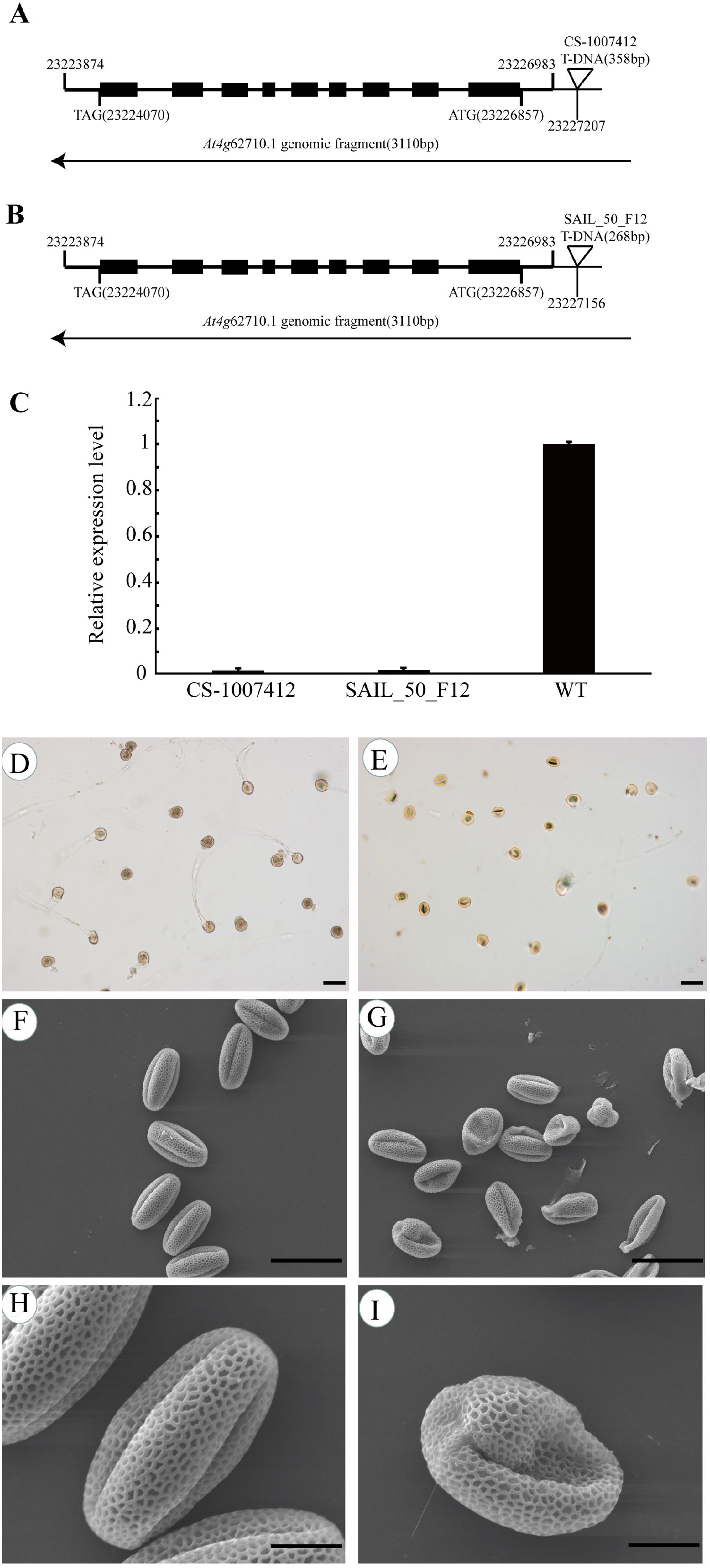
The phenotype of Arabidopsis *βvpe* mutant plants (A) CS_1007412 T-DNA insertion positions in *At1g62710.1*. (B) SAIL_50_F12 T-DNA insertion positions in *At1g62710.1*. (C) *βVPE* expression analyses in mutants. (D) Germination rate of wild-type pollen. (E) Germination rate of mutant pollen. (F) and (G) Scanning electron microscopy of wild-type pollen. (H) and (I) Scanning electron microscopy of mutant pollen. (D), (E), (F) and (G) bar = 50 um; (H) and (I) bar = 10 um.

### 3.3 Anther development in βvpe mutants

Both wild-type and *βvpe* anthers were examined to further clarify the pollen development process in the *βvpe* mutant. At stage 10, no obvious differences were observed between the wild type and the *βvpe* mutant (Figure 3A, B, E, and F). At stage 11, well-developed microspores were expanded in wild-type anthers. In contrast, some microspores were still vacuolated in the *βvpe* mutant (Figure 3C and G). At stage 13, mature pollen grains were formed in the wild type. However, although some *βvpe* pollen grains exhibited normal development, some pollen grains were shrunken and defective (Figure 3D and H). These results indicate that pollen development was partially abnormal and the number of mature pollen grains was significantly decreased in *βvpe* mutants.

**Figure 3.**
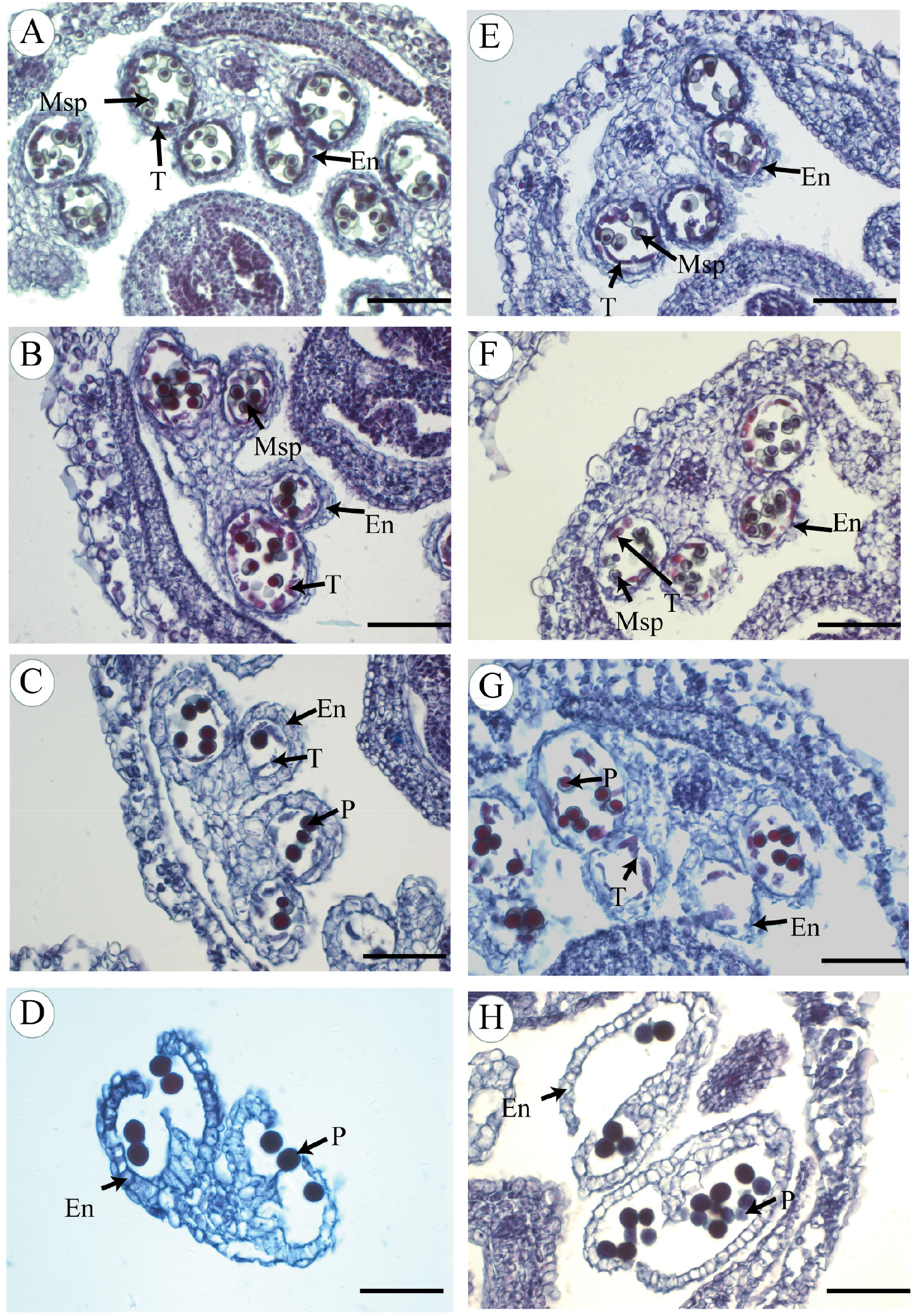
Anther development in the wild type and *βvpe* mutant The wild type (A-D) and cep1 mutant (E-H) during anther development. (A) and (E) stage 8; (B) and (F) stage 10; (C) and (G) stage 11; (D) and (H) stage 13. Bar = 50 um. dP, degenerated pollen; En, endothecium; Msp, microspore; P, pollen; T, tapetum.

### 3.4 Abnormal pollen development in βvpe mutants

We performed TEM to investigate pollen development in *βvpe* mutants. No obvious differences were observed in microspore development between *βvpe* mutants and the wild type before stage 10. At stage 9, the development of the orderly microspore exine structure proceeded in both the wild type and in *βvpe* mutants (Figure 4A, F, and K). At stage 10, many oil bodies were present in the microspore, and one nucleus was present in the lenticular-shaped generative cell in the wild type. In *βvpe* mutants, unlike in the wild type, pollen cytoplasm development was incomplete, with few oil bodies and an indistinct generative cell. Some of the pollen grains were shrunken and abnormally shaped in *βvpe* mutants at this stage (Figure 4B, G, and L). At stage 11, the immature pollens grains continued to develop and the vacuole completely disappeared in the wild type (Figure 3C). However, numerous small vacuoles were distributed throughout the cytoplasm of the *βvpe* pollen grains (Figure 4C, H, and M). At stage 12, the typical pollen wall was completely established in the wild type. In the mutants, immature pollen still contained numerous small vacuoles and some of the pollen grains were shrunken (Figure 4D, I, and N). At stage 13, wild-type pollen was fully mature. In the *βvpe* mutants, the abnormal pollen grains were shriveling (Figure 4E, J, and O). These results indicated that pollen development was not normal in *βvpe* mutants. The stage of vacuole breakage was significantly delayed and the shriveling and invagination of some of the pollen grains indicated that they were not mature in *βvpe* mutants. These results indicate that pollen development in *βvpe* mutants was impaired, resulting in delayed vacuolar degradation and collapsed pollen grains.

**Figure 4.**
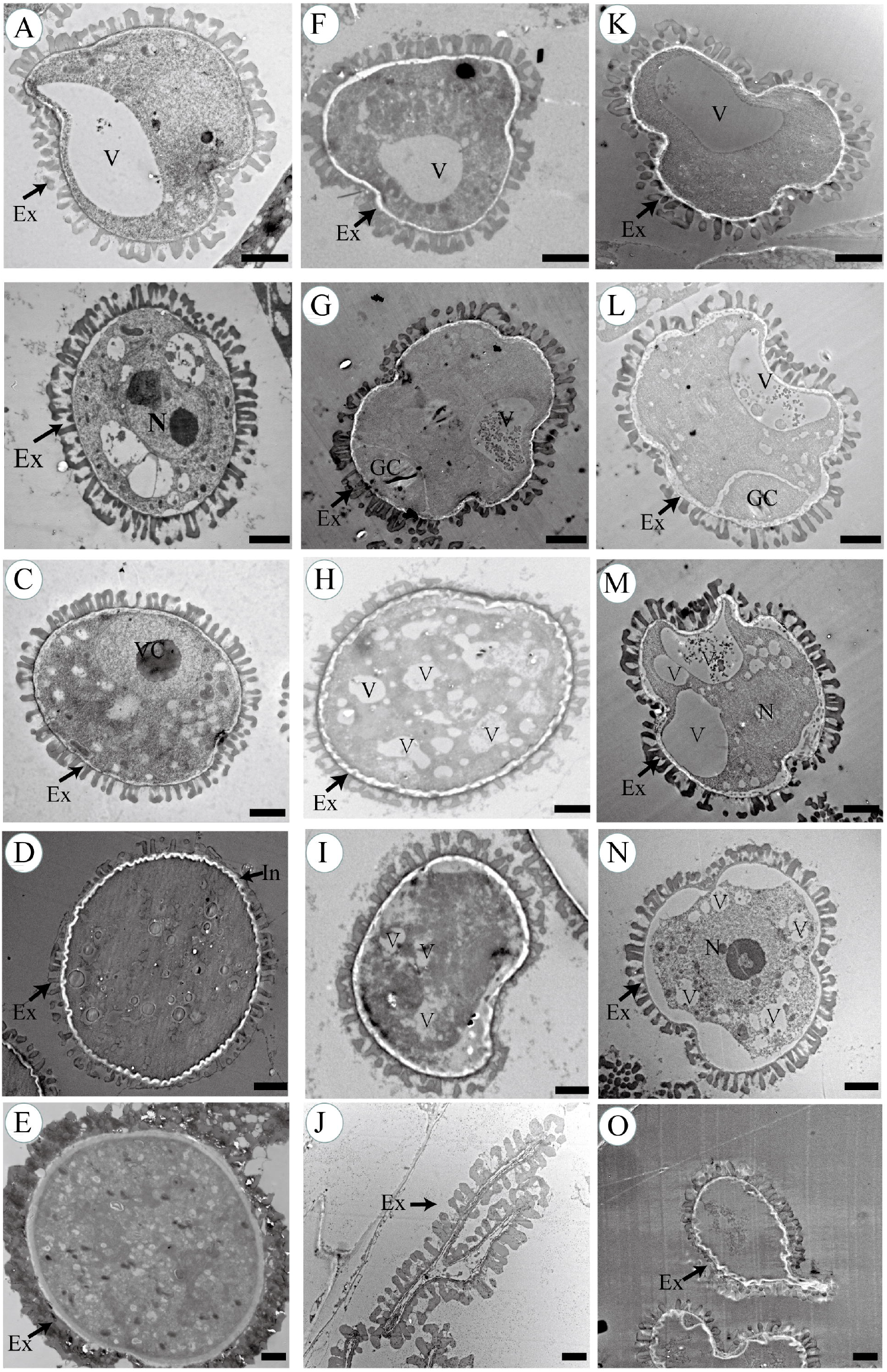
Transmission electron micrographs of microspores from the wild type and *βvpe* mutant Microspores of different developmental stages in the wild type (A-E), *βvpe* mutant CS_1007412 (F-J), and *βvpe* mutant SAIL_50_F12 (K to O): (A), (F), and (K) stage 9; (B), (G), and (L) stage 10; (C), (H), and (M) stage 11; (D), (I) and (N) stage 12; (E), (J), and (O) stage 13. Bar = 2 mm. Ex, exine; GC, generative cell; In, intine; N, nucleus; T, tapetal cell; V, vacuole; VC, vegetative cell.

### 3.5 Abnormal degradation of the tapetum in βvpe mutants

A transmission electron microscopy (TEM) assay was used to visualize differences in the tapetal cells between the wild type and *βvpe* mutants in stages 9-12. At stage 9 in the wild type, the tapetal cell wall had completely degraded and the tapetosome containing many lipid materials appeared in the binucleate tapetal cell (Figure 5A). However, the tapetal cell wall partially remained in the *βvpe* mutant, indicating that the mutant’s tapetal cells had failed to transform properly into the polar secretory type and had only formed a few secretory vacuoles and vesicles (Figure 5E and I). At stage 10, the nuclei of wild-type tapetal cells had already degraded, the numbers of tapetosomes and elaioplasts were enriched, and the tapetal cells continuously released fibrillar materials into the anther locules (Figure 5B). In contrast, few elaioplasts and almost no tapetosomes had formed in the *βvpe* mutant, resulting in few fibrillar materials being released from its tapetal cells (Figure 5F and J). At stage 11, wild-type tapetal cells reached the end of their PCD and were filled with tapetosomes and elaioplasts (Figure 5C). However, in the *βvpe* mutant, tapetal cells wall still remained, and while the cytoplasm of tapetal cells had undergone degeneration, no obvious tapetosomes or elaioplasts were present (Figure 5G and K). At stage 12, tapetal cell degeneration had finished in the wild type (Figure 5D), but tapetal cell remnants with partially undegenerated cell walls remained in the *βvpe* mutant (Figure 5H and L). These results indicate that the degradation of tapetal cells in *βvpe* mutants was abnormal and that the formation of secretory organelles had largely decreased.

**Figure 5.**
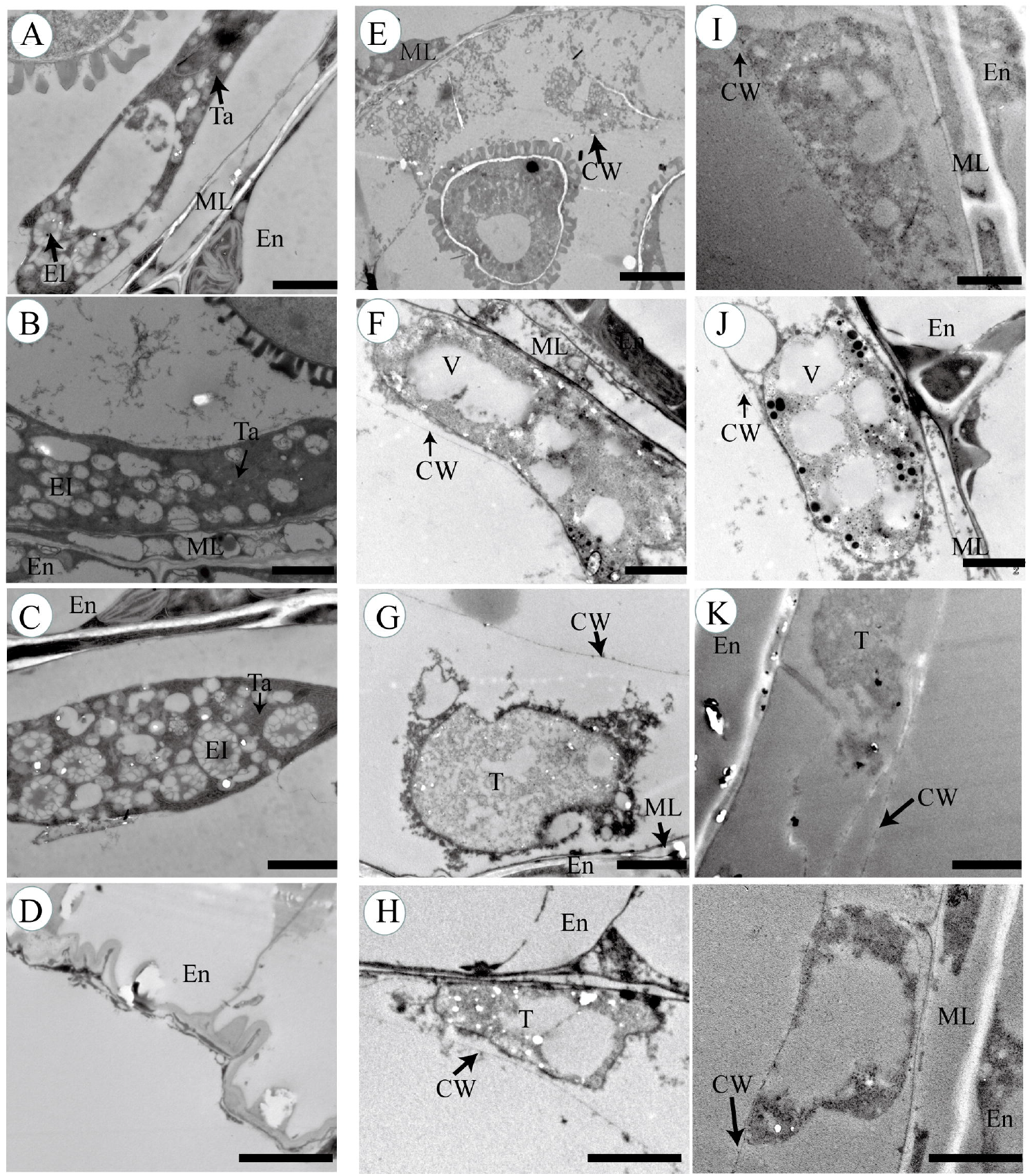
Transmission electron micrographs of anthers from the wild type and *βvpe* mutant (A)-(D) Wild type; (E)-(H) *βvpe* mutant CS_1007412; (I)-(L) *βvpe* mutant SAIL_50_F12: (A), (E), and (I) stage 9; (B), (F), and (J) stage 10; (C), (G), and (K) stage 11; (D), (H), and (L) stage 12. Bar = 2 mm. CW, cell wall; El, elaioplast; En, endothecium; ML, middle layer; T, tapetal cell; Ta, tapetosome; V, vacuole.

### 3.6 Proβvpe: βvpe translational fusion complements the βvpe mutation

A complementation experiment was performed to confirm that the *βvpe* mutant phenotype was attributable to the loss of *βVPE* function. An 1847-bp promoter of *βVPE* (Pro*βVPE*) and the 1713-bp *βVPE* cDNA sequence were cloned into the pCAMBIA 1300 vector and introduced into *βvpe* mutant plants. Among 46 independent transgenic plants that grew to maturity, 35 plants had restored normal fertility, 8 were partially fertile, and the remaining 3 plants were similar to the *βvpe* mutant, with no restoration of fertility (Figure 7). The negative control lines transformed with the empty pCAMBIA 1300 vector revealed a sterile phenotype similar to that of the *βvpe* mutant. The 35 restored transgenic lines displayed normal tapetal degeneration and fertile pollen grains.

**Figure 6.**
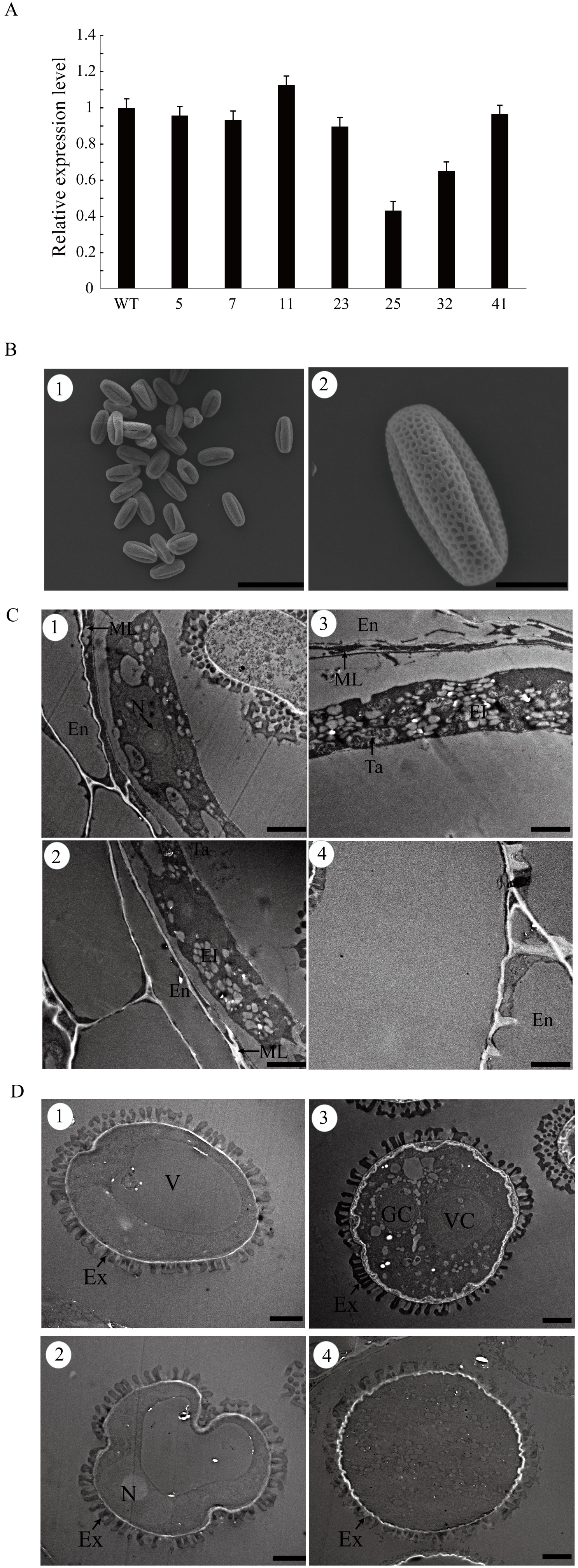
Complementation of the *βvpe* mutant by *βVPE* cDNA (A) qRT-PCR of *βVPE* expression in bud tissues of complementation lines. (B) Scanning electron microscopy of mature pollen grains of complementation lines: (B1) bar = 50 um; (B2) bar = 10 um. (C) Transmission electron micrographs of the tapetum development in complementation lines: bar = 2 mm. (D) Transmission electron micrographs of microspore development in complementation lines: bar = 2 mm.

The restored transgenic lines displayed normal tapetal degeneration, pollen development, and fertile pollen grains. The mature pollen grains of the restored transgenic lines and the wild type were uniformly spheroid and had finely reticulate ornamentation on their surfaces (Figure 7B). The tapetum degradation and microspore development of anthers during stages 9-12 of the restored transgenic lines were consistent with wild-type development. The formation of tapetosomes and elaioplasts was normal and the degradation of the cell wall in tapetal cells was complete in the restored transgenic lines (Figure 7C). The development of pollen in the restored transgenic lines was also the same as that of the wild type (Figure 7D). These results indicate that the phenotype of the *βvpe* mutant is caused by the loss of βVPE.

**Figure 7.**
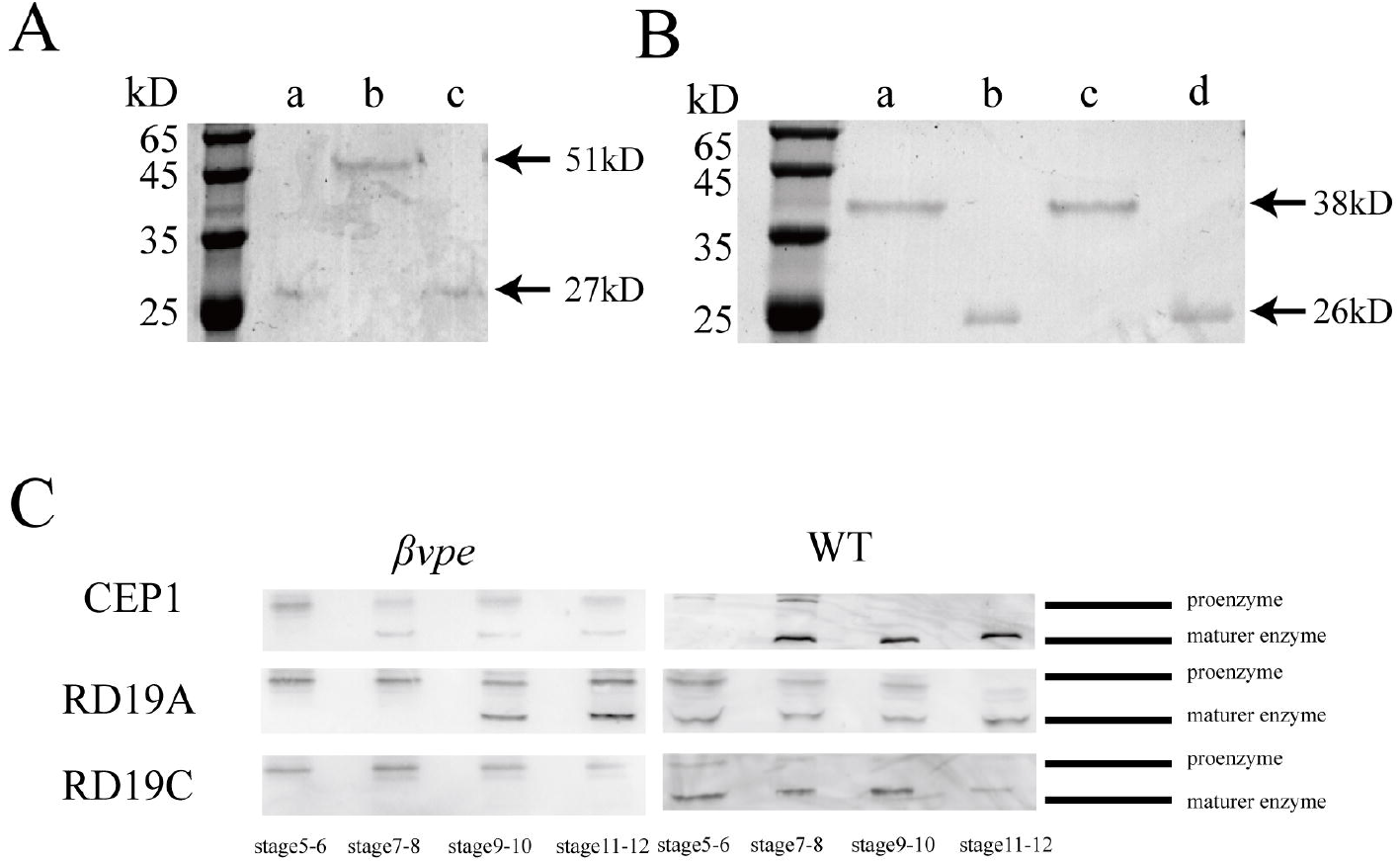
Activation of proproteases CEP1, RD19A, and RD19C (A) Purified and mature recombinant βVPE: (a) and (c) mature βVPE protein created by self-cleaving at pH 5.2; (b) recombinant pro-βVPE. (B) Purified and mature recombinant CEP1: (a) recombinant pro-CEP1 at pH 7.0; (b) mature CEP1 protein created by self-cleaving at pH 3.0; (c) pro-CEP1 at pH 5.2; (d) mature CEP1 protein created by mediation of βVPE. (C) Immunoblot analysis of total anther protein extracts from stages 5 to 12 with a ntimatu re-CEP 1 antibody, antimature-RD19A antibody, and antimature-RD19C antibody.

Quantitative RT-PCR (qRT-PCR) analysis of *βVPE* transcript levels in the transgenic plants revealed that the transgenic lines with restored fertility contained similar numbers of transcripts as the wild type (Figure 6A).

### 3.7 βVPE can transform the precursors CEP1, RD19A, and RD19C to mature proteins *in vivo*

To further characterize the properties of βVPE, induction of the expression construct pET30a-βVPE, which encodes the full-length βVPE cDNA minus the signal peptide, was performed in *Escherichia coli*, resulting in overexpression of the recombinant protein. The molecular weight of the purified recombinant proβVPE was 51 kD according to SDS-PAGE analysis (Figure 7A, lanes 1 and 3). The mature protein had a molecular weight of 27 kD and self-cleaved from the proβVPE protein at pH 5.2 (Figure 7A, lane 2).

To further characterize whether VPE could transform the maturation of CEP1 *in vitro*, we induced the expression construct pET30a-CEP1, which encodes the full-length CEP1 cDNA minus the signal peptide, in *Escherichia coli*, resulting in overexpression of the recombinant protein. The molecular weight of the purified recombinant CEP1 was 38 kD, according to SDS-PAGE analysis, and it transformed to the 26-kD mature protein by self-cleaving at pH 5.2 (Figure 7B, lanes 1 and 2). The CEP1 proprotein was not able to transform to the mature protein by self-cleaving at pH 5.2 (Figure 7B, lanes 3). When mature VPE was added, the 26-kD mature protein CEP1 was detected at pH 5.2 (Figure 7B, lanes 4).

To determine whether VPE could transform the inactive precursor proteases to mature proteins *in vivo*, immunoblotting was employed to investigate three cysteine proteases (CEP1, RD19A, and RD19C) from stages 5-12 in both wild-type and *βvpe* mutant anthers. In the wild type, the 38-kD CEP1 proenzyme appeared in stages 5-6, the mature 26-kD enzyme appeared during stages 7-8, and only the mature 26-kD enzyme was found abundantly in the anther during stages 9-12. However, in *βvpe* mutants, the content of mature 26-kD CEP1 was much lower, and a large amount of proenzyme still existed during stages 9-12 (Figure 7C). These results indicate that self-cleavage at pH 3.0 and the catalysis of βVPE in acidification of the vacuole work together to produce mature CEP1 in anthers.

In the wild type, the RD19A proenzyme was present in stages 5-10, but the 26-kD mature enzyme did not appear until stages 7-8, and only the mature 26-kD enzyme was found abundantly in the anther during stages 11-12. However, in the *βvpe* mutant, the content of mature RD19A enzyme was greatly diminished, the mature enzyme appeared during stages 9-10, and considerable quantities of RD19A proenzyme were still observed during stages 9-12 (Figure 7C). These results indicate that RD19A proenzyme did not self-cleave in response to acidification of the vacuole when βVPE was not present, but that it was able to transform to mature RD19A enzyme after vacuole rupture without the presence of βVPE (Figure 7C).

In the wild type, the RD19C proenzyme was present in stages 5-6, and the mature 26-kD mature enzyme appeared during stages 7-12. However, only the RD19C proenzyme was present during stages 5-12 in the *βvpe* mutant (Figure 7C). These results indicate that βVPE processing is the only route for the maturation of RD19C in the anther.

Together, these results demonstrate that βVPE is important for the transformation of the proproteases CEP1, RD19A, and RD19C into fully active mature enzymes that are necessary for proteolytic processing during anther development.

## 4. Discussion

### 4.1 βVPE should transform to its mature form by self-cleavage in response to acidification of the vacuole during anther development

Vacuolar processing enzymes are endopeptidases with substrate specificity to asparagine residues. They are synthesized as inactive larger proprotein precursors, from which the C-terminal and N-terminal propeptides are sequentially self-catalytically removed to produce the active mature forms under acidic conditions. Previous research has shown that the 56-kD pro-γVPE is self-catalytically converted to a 43-kD intermediate form and then to the 40-kD mature form at pH 5.5 (Kuroyanagi *et al.*, 2002). In Arabidopsis seeds, βVPE has a 37-kD intermediate form and a 27-kD mature form (Shimada *et al.*, 2003). In our research, high levels of expression of βVPE were detected in stage 5-8 Arabidopsis anthers. βVPE existed as pro-βVPE at stages 5-6 and was self-catalytically converted to the 27-kD mature form in stages 5-8 *in vivo*. A further assay using prokaryotic expression *in vitro* revealed that pro-βVPE transformed to mature protein at pH 5.2. Previous studies have reported that vacuole acidification begins during late stage 6 and vacuole degradation is complete by late stage 8. Pro-βVPE should transform to its mature form by self-cleaving during the early stages of vacuole acidification in anther development. With vacuolar rupture complete by late stage 8, mature βVPE is released into the cytoplasm, where it is degraded. This finding suggests that βVPE acts primarily in vacuoles, not in the cytoplasm, during anther development. Tapetal development is controlled by a complex transcriptional regulatory network. Many transcription factors are involved in anther cell differentiation and tapetal development. An analysis of the known microarray data revealed that βVPE expression is downregulated 1.3-fold, 4.0-fold, and 3.9-fold in ms1, ems1, and spl mutants, respectively, and upregulated 3.6-fold in tdf1 mutants (Vizcay-Barrena *et al.*, 2006; Wijeratne *et al.*, 2007; Phan *et al.*, 2011). However, βVPE expression is not significantly altered in the dyt1, rpk2, ams, ashh2, mia, myb80, and roxy1roxy2 mutants, whereas other papain-like cysteine proteases, such as RD19C, RDL1, RD19A, THI1, and RD21A, show changes in expression level to varying degrees (Bernoux *et al.*, 2008; Feng *et al.*, 2012; Ito *et al.*, 2007; Li *et al.*, 2017; Phan *et al.*, 2011, 2012; Yang *et al.*, 2007). SPL/NZZ and EMS1/EXS are expressed during anther differentiation and probably act as early regulators of βVPE expression around stage 5. These results suggest that βVPE is involved in the SPL/NZZ, EMS1/EXS, and TDF1 pathways regulating tapetal development and degeneration.

### 4.2 βVPE directly participates in the maturation of cysteine proteases

Previous reports have shown that premature protease is transported to the protease vesicles, presumably via the endoplasmic reticulum, and then transported into the vacuole, ricinosomes, or lysosome, and transformed to mature protease in response to acidification at stage 6 by self-cleavage or protease-dependent maturation. For example, castor bean CysEP and tomato CysEP are active at pH 4-6.5 (Senatore *et al.*, 2009; Greenwood *et al.*, 2005). Previous studies in our lab have shown that the proenzyme of CEP1 is transformed to the mature form by self-hydrolysis in vacuoles at pH 3.0 during Arabidopsis anther development (Zhang *et al.*, 2014).

Our results indicate that the mature βVPE enzyme could activate pro-CEP1 *in vitro* and transform the inactive precursors CEP1, RD19A, and RD19C to mature proteins *in vivo*. The maturation of CEP1 and RD19A was seriously suppressed, and the mature form of RD19C completely undetectable, in the *βvpe* mutant. Previous research in our lab revealed that pro-CEP1 is transformed to its mature form by self-hydrolysis at stages 6-8 with acidification of the vacuole but before vacuole rupture. In the *βvpe* mutant, however, CEP1 proenzyme was detected during stages 9-13. This finding suggests that the CEP1 proenzyme transforms in two ways: by self-hydrolysis and by the action of βVPE. Self-hydrolysis of βVPE occurs at pH 5.2 and self-hydrolysis of CEP1 occurs at pH 3.0, suggesting that the maturation time of VPE is earlier than that of CEP, and that βVPE could activate the transformation of CEP1 into a mature protein in vacuoles. The maturation of RD19A was also seriously suppressed, suggesting that βVPE processing is one way in which proRD19A is transformed to a mature protein. However, RD19A proenzyme transformed to mature RD19A enzyme after vacuole rupture without the mediation of βVPE. The mature form of RD19C was completely undetectable in the *βvpe* mutant, suggesting that the transformation of proRD19C to a mature protein can only occur through a βVPE-mediated process.

These results indicate that βVPE is involved in the maturation of cysteine proteases during tapetum development.

### 4.3 βVPE is indirectly involved in pollen development and tapetal cell degradation

Many plant cysteine proteases are implicated in a variety of PCD events in various tissues, including xylogenesis (Han *et al.*, 2012), leaf and flower senescence (Chen *et al.*, 2002; Senatore *et al.*, 2009), seed development and germination (Greenwood *et al.*, 2005), and tapetum development (Zhang *et al.*, 2014). The tapetum plays an important role in microspore development, providing enzymes for the release of microspores from four bodies as well as nutrients for pollen development and essential components for the development of pollen walls. The premature termination of PCD in tapetal cells can interrupt the nutrient supply for microspore development, eventually leading to male infertility (Ko *et al.*, 2003; Varnier *et al.*, 2005). Previous research has shown that the cysteine protease CEP1 is expressed specifically in the tapetum, and that tapetal PCD is aborted and pollen fertility is decreased with abnormal pollen exine in the *cep1* mutant (Zhang *et al.*, 2014). In our study, loss of βVPE function in the *βvpe* mutant caused the failure of some CEP1 proproteins to transform to mature enzymes. A TEM analysis confirmed the failure of tapetal cell wall degeneration, a large decrease in the formation of elaioplasts and tapetosomes, and abnormal pollen development in the *βvpe* mutant in stages 9-12 of anther development. Because βVPE is degraded at stages 9-12 of anther development, it seems that βVPE participates indirectly in pollen development and tapetal cell degradation. The main reason for abnormal pollen development and failure of tapetal cell wall degeneration in the *βvpe* mutant may be due to the failure of other proproteases, such as CEP1, RD19A, and RD19C, to mature normally. Our results also showed that some CEP1 proproteases transformed to mature protein even in the absence of βVPE, likely by self-hydrolysis. This may explain the presence of some normal pollen grains (50%) in *βvpe* mutants.

In conclusion, during the early stages of anther development βVPE triggers the activation of other proproteases, such as CEP1, RD19A, and RD19C, to transform into mature proteins. First, pro-βVPE transforms to the mature βVPE enzyme in vacuoles. Then, βVPE activates the proproteases before vacuole rupture. After vacuole rupture, the mature βVPE enzyme quickly degrades. In our research, we first discovered that βVPE can activate proproteases. Our findings provide valuable insights into the regulatory network determining protein levels in the tapetum and in identifying the involvement of βVPE in anther development.

## Acknowledgements

This work was funded by the Fundamental Research Funds for the Central Universities (No. BLX2015-37) and Chinese National Science Fund (No.31570582).

The English in this document has been checked by at least two professional editors, both native speakers of English. For a certificate, please see:

http://www.textcheck.com/certificate/NxnrqK

